# Fast Optical Sectioning for Widefield Fluorescence Mesoscopy with the Mesolens based on HiLo Microscopy

**DOI:** 10.1101/374884

**Authors:** Jan Schniete, Aimee Franssen, John Dempster, Trevor Bushell, William Bradshaw Amos, Gail McConnell

## Abstract

We present here a fast optical sectioning method for optical mesoscopy based on HiLo microscopy, which makes possible imaging of specimens of up to 4.4 mm × 3 mm × 3 mm in volume in under 17 hours (estimated for a z-stack comprising 1000 images excluding computation time) with subcellular resolution throughout. Widefield epifluorescence imaging is performed with the Mesolens using a high pixel-number camera capable of sensor-shifting to generate a 259.5 Megapixel image, and we have developed custom software to perform HiLo processing of the very large datasets. Using this method, we obtain comparable sectioning strength to confocal laser scanning microscopy (CLSM), with sections as thin as 6.8±0.2 μm and raw acquisition speed of 1 minute per slice which is up to 30 times faster than CLSM on the full field of view (FOV) of the Mesolens of 4.4 mm with lateral resolution of 0.7 μm and axial resolution of 7 μm. We have applied this HiLo mesoscopy method to image fixed and fluorescently stained hippocampal neuronal specimens and a 5-day old zebrafish larva.

## Introduction

The Mesolens is a novel microscope objective lens that combines a high numerical aperture (NA) of 0.47 with a large field of view (FOV) of up to 6 mm. It has been used as the basis for CLSM to generate high quality optical sections of large specimens such as e12.5 mouse embryos with 700 nm lateral and 7μm axial resolution ^1^. The combination of large field of view and high spatial resolution means that confocal images of around 20,000 pixels × 20,000 pixels are needed for Nyquist sampling. Even with a short pixel dwell time of 1 μs confocal imaging is slow, taking around 400 seconds per image. Z-stacks of large volume specimens can take from several hours up to days or even weeks to acquire, depending on the total volume, number of channels used, and any frame averaging to improve the signal-to-noise ratio. Although the Mesolens has proven utility in biomedicine ^1,2^, a fast widefield optical sectioning method is essential to reduce acquisition time to make the Mesolens suitable for rapid high-resolution imaging of large volume specimens.

Widefield techniques capable of performing optical sectioning are highly sought after in biological imaging to reduce photodamage and photobleaching as well as increasing signal to noise ratio (SNR) at high acquisition speed. Each method has its own strengths and weaknesses and the choice depends on the specimen of interest.

Widefield two-photon microscopy (W2PM) has been reported for *in vivo* imaging ^3^ and it has been shown to produce less photobleaching than single-photon excitation without the need to scan the beam over the field of view, capable of reaching 100Hz acquisition speed ^4^. Furthermore, W2PM can perform optical sectioning when the peak intensity overcomes the threshold for 2P absorption in the focal plane only using temporal focusing ^5,6^. On the downside, the 2P absorption process is non-linear and requires ultra-short laser pulses on the order of few hundred femtoseconds. W2PM would require very high peak intensity radiation propagating through the Mesolens to overcome the threshold to excite fluorescent molecules at the sample, potentially damaging the optical elements.

Selective plane illumination microscopy (SPIM), also called light sheet microscopy (LSM) has received a lot of attention in recent years. Since its original conception in 1902 ^7^, SPIM has been reintroduced for modern microscopy techniques ^8^. The biggest advantage is the lack of illumination outside the focal plane. This leads to minimal phototoxicity and photobleaching. However, the side-on illumination can lead to inhomogeneous brightness across the field of view and scattering samples can cast ‘shadows’ in the lateral direction. Techniques have evolved to counter these effects, e.g. dual side illumination ^9^ and the use of non-diffracting beams allow the creation of thin light sheets over a comparably large FOV up to 1 mm ^10^. A tenfold increase in light sheet FOV coverage over standard Gaussian light sheets has been reported ^11^, but the total volume is still ~10 times less than can be accommodated with the Mesolens at similar spatial resolution (based on a CFI S Fluor 10x objective lens with 0.5 NA and 1.2 mm working distance). For the Mesolens, because of its unique combination of high NA and large FOV, SPIM is not capable of generating a light sheet that has a long enough Rayleigh length to cover the full field of view at ~7μm thickness. The only method to cover such a large FOV with a thin sheet would be to scan the light sheet and acquire multiple images, similar to stitching and tiling on standard microscope lenses but with the FOV remaining unchanged and the illumination moving through several positions in the FOV. This method would defeat the purpose of using the Mesolens and greatly increase acquisition time.

Structured illumination microscopy (SIM) is a super-resolution optical imaging technique that inherently provides optical sectioning as it only regards modulated, in-focus signal for the final image ^12,13^. A related method called HiLo microscopy has been developed recently ^14^ which makes use of the optical sectioning capability of SIM without any super-resolution content. Only two images are required for HiLo microscopy as opposed to 4 or more images for SIM techniques, which can reduce phototoxicity and photobleaching effects ^15,16^. HiLo uses post processing of one structured and one uniform illumination image to achieve the result. The optical sectioning strength of HiLo has been reported to be comparable to CLSM and in principle is only limited by the camera exposure time ^17^. The HiLo method can be implemented inexpensively by using a diffuser to create laser speckle with coherent (laser) light or with a grating and incoherent light. This diffuser-based method of generating laser speckle has been reported to be more robust than using a grating to study scattering samples ^17^ and the contrast of speckle can be easily adjusted to suit different samples ^18^. SIM does provide super-resolution but it is not straightfoward to implement with the Mesolens: structured illumination would have to be introduced in the back aperture plane of the condenser because the back focal plane of the objective is not accessible in the current Mesolens design. As a direct result, thick samples could not be imaged because the grating contrast would quickly deteriorate because of scattering.

We therefore chose to implement HiLo microscopy in transmission illumination with laser speckle illumination with the Mesolens to obtain optical sections and investigated sectioning strength, quality and speed compared to CLSM.

## Results

It was found that the lowest setting for the optical sectioning parameter σ where the dependence of optical section thickness on σ was still linear corresponded to a frequency of 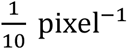. The scaling in the MATLAB script was therefore adjusted to have this value as a minimum when σ was set to 1. At this setting an optical section thickness of 6.8±0.2 μm (mean ± standard deviation of five measurements) was measured by evaluating the average FWHM of Gaussian fits to intensity line plots through the processed image of the tilted fluorescent layer discussed in the Materials and Methods section. Similarly, the minimum optical section thickness of the ImageJ plugin attainable was measured at 6.6±0.3 μm. **Figure 1** shows the tilted layer processed with our MATLAB script compared to the HiLo ImageJ plugin and optical section thickness measurements at σ settings ranging from 1 to 10 for the plugin and MATLAB script respectively. The range of σ was chosen such that the resulting section thickness scaling and range was comparable between plugin and MATLAB script.

**Figure 1:**
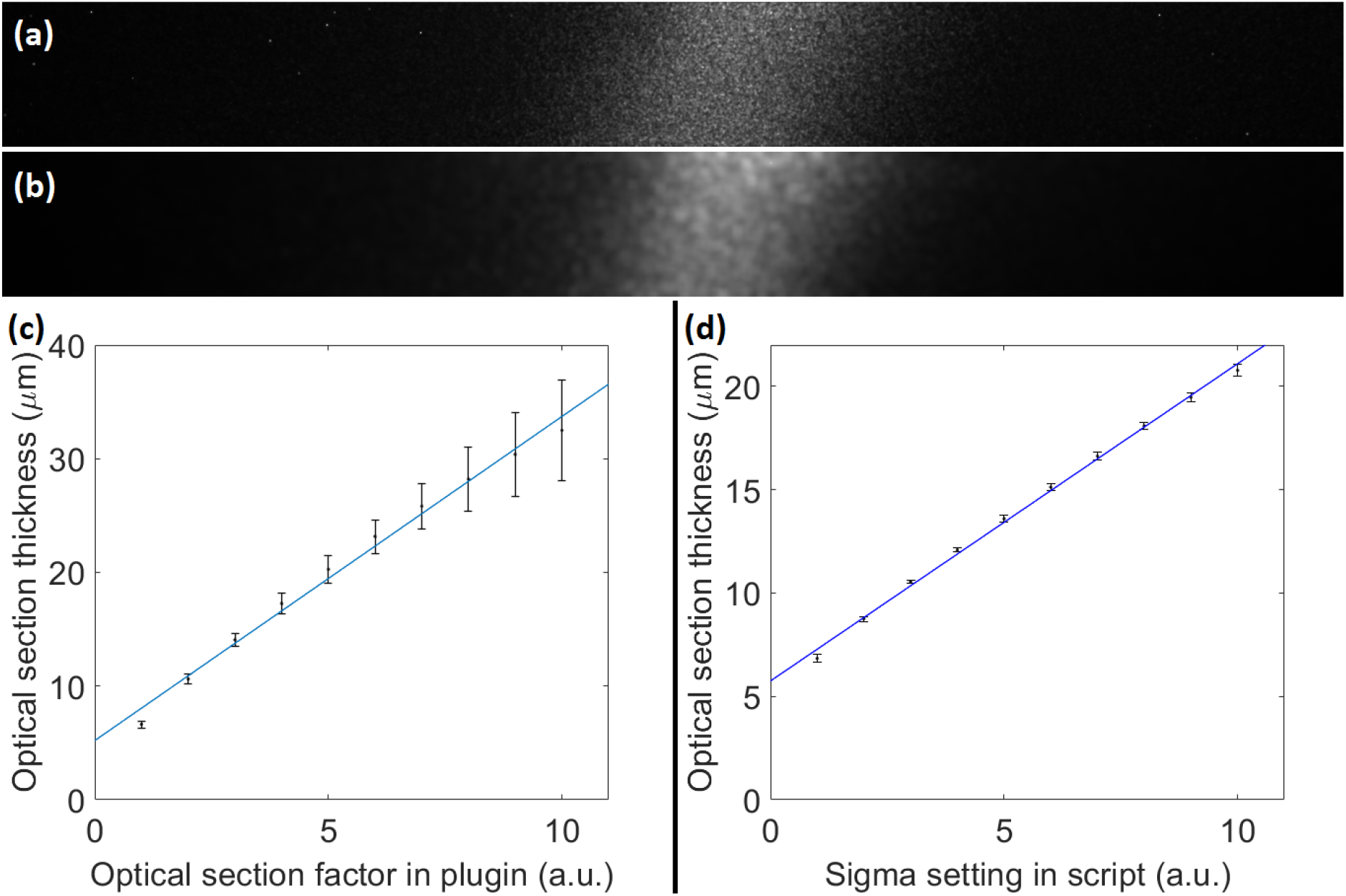
Comparison of optical sectioning strength. A tilted layer of fluorescent dye was processed with the HiLo ImageJ plugin and the our MATLAB script. An example of the HiLo processed layer is shown for (a) the plugin with the optical sectioning scaling parameter σ set to 2 pixel-1 and (b) the MATLAB script with σ set to 3 pixel-1 which corresponded to approximately the same optical section thickness. The thickness as a function of σ was obtained by processing the same data of the tilted fluorescent layer with the plugin (c) and our MATLAB script (d) and calculating the average full-width-half-maximum (FWHM) of Gaussian fits to five horizontal intensity line plots through the processed images. The two processing modalities reached a minimum section thickness of 6.6±0.3 μm (plugin) and 6.8±0.2 μm (MATLAB) at the lowest setting for σ.

As expected from other work in the microscopic domain ^18^, it was found that the speckle pattern for the structured illumination image needed to be coarser when imaging thicker samples (more than ~100 μm thick). In the ideal case, the transverse size of an imaged speckle grain was determined by the illumination NA ^14,17^. However, for thicker samples the grain size was increased to approximately 20 pixels (~5 μm at 9× chip shifting) to maintain high contrast in the difference image for the in-focus regions of the image. The choice of the σ parameter had to be made according to the coarseness of said pattern such that several imaged grains would fit in the sampling window for contrast evaluation so no artefacts of speckle structure translated through to the final image. Specimens of fixed and fluorescently stained hippocampal mouse neurons and a 5-day-old zebrafish larva specimen were imaged with the Mesolens and processed in MATLAB using suitable σ settings to evaluate performance of the method with thick and uncleared biological specimens.

The ~150 μm thick zebrafish stained with Acridine Orange was imaged and HiLo processed in MATLAB with σ=2. This was the lowest value for the combination of sample thickness and choice of speckle coarseness where no artefacts were observed in the final image. The excitation wavelength was 488 nm with ~4 mW laser power at the sample. The fluorescence signal was filtered at 540 nm with spectral full-width-half-maximum of ~20 nm and exposure time of the camera detector was 400 ms for each of the nine sensor positions. Supplementary Supplementary Video **1** shows the data as a z-series video, with the z-stack comprising 61 images taken with 3 μm steps in axial position. The FOV was cropped to roughly the size of the zebrafish (3.2 mm × 1.2 mm). The zoomed-in eye shows improved contrast. Individual cells are clearly visible as well as the broader structure of the surrounding tissue. **Figure 2** shows a transmission illumination widefield image (the uniform image also used for processing) compared to an optical section from the 61 image z-stack at approximately 100 μm depth in the sample. An improvement in signal to noise ratio was evident across the whole field of view and zoomed-in regions of interest revealed fine detail that was not clearly visible in the widefield image.

**Figure 2:**
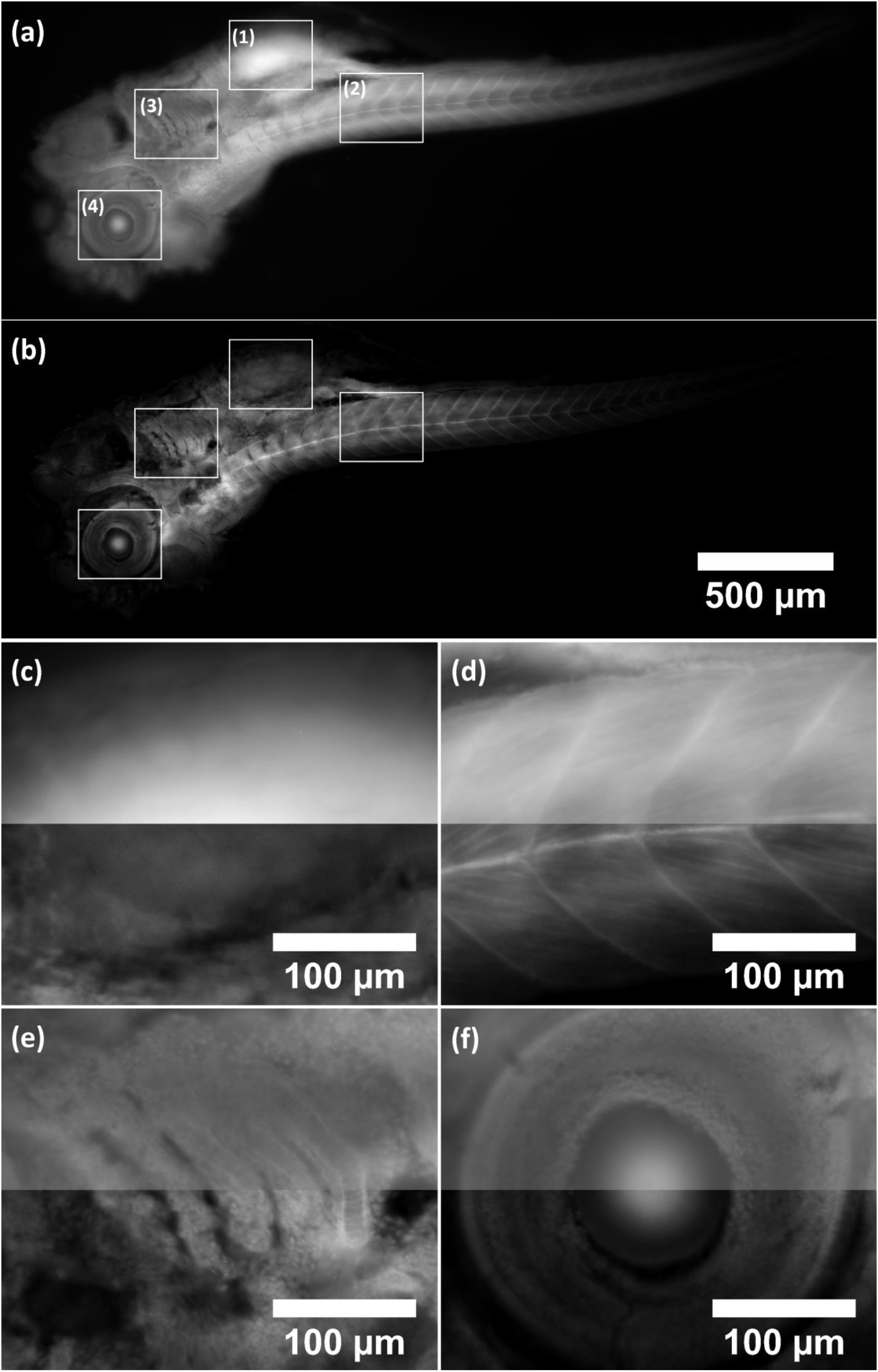
Standard widefield image (a) compared to Mesolens HiLo-processed optical section (b) of zebrafish stained with Acridine Orange. Insets (c-f) show zoomed-in regions of interest (1-4) where the widefield and HiLo image were merged together (top half was widefield, bottom half was HiLo image). Contrast improvement was evident across the whole image and regions of interest showed fine detail that was barely noticeable in the widefield image. Optical sectioning parameter σ was set to 2, corresponding to ~8.7±0.1 μm. With σ set to 1 there were too many artifacts. There was still a hint of inhomogenous brightness with σ=2 but not so severe that false detail emerged in the final image. Setting σ higher would have resulted in an unnecessarily thick section.

For sparse samples such as neurons, the sectioning parameter σ was set to 1. This was the lowest setting at which the sectioning curve in **Figure 1**.behaved linearly and was chosen to obtain the thinnest sections without speckle artefacts in the final image. Supplementary **Supplementary Video 2** shows fluorescently stained mouse hippocampal neurons sectioned at this setting. The excitation wavelength, detection bandpass average power at the sample were the same as for the zebrafish specimen. The camera exposure setting was 200 ms (for each sensor position). Optical sectioning revealed the axial extent of dendrites and cell bodies (soma) and improved contrast to clearly show the unstained centre of the soma where the nucleus is located. Acquiring the data took ~27 minutes per stack (uniform and speckle illumination) and another ~26 minutes to process in MATLAB.

Acquisition of one image pair (speckle and uniform illumination) took approximately 1 minute for full 4.4 mm FOV and Nyquist sampled images (9× chip-shifting). It was found that the frame time, i.e. the time it took to acquire an image and start the next acquisition, was mainly taken up by transferring the image from the camera to the PC and temporarily saving it on the hard drive. Hence the actual exposure did not significantly impact the time to acquire a whole z-stack. The 25-image z-stack of neurons took ~27 minutes to acquire per illumination at 200 ms camera exposure. Processing a full FOV full resolution image-stack in MATLAB took ~1 minute per image pair. Image processing was always performed post acquisition and did not increase imaging time. Furthermore, the same raw images could be processed at different settings without the need to re-acquire data. Compared to CLSM this method acquires raw data ~30 times faster, including processing it is still ~15 times faster, estimated based on Nyquist sampled CLSM full FOV image with three times averaging (~30 minutes acquisition).

## Discussion

We have shown here a fast widefield optical sectioning method for the Mesolens using HiLo microscopy capable of section thickness of 6.5±0.2 μm over the full FOV of 4.4 mm, on par with CLSM on the Mesolens which can generate sections of around 5μm thickness. The lateral resolution remains unaffected by HiLo microscopy because the high spatial frequency, i.e. high resolution, content of the final image is obtained directly by high-pass filtering the uniform image. Since the uniform image is a standard widefield camera image, no fine detail is lost ^18^. This method presents a significant speed advantage over CLSM at comparable sectioning strength of ~5μm, being 30 times faster in raw data acquisition. We elected to write our own script in MATLAB rather than use the existing ImageJ HiLo plugin. Although the plugin performed very well when we initially tested it with small datasets, it was very slow (~16 times slower than the current MATLAB implementation) when processing large Mesolens data and in some cases data could not be processed at all due to memory limitations (even when using a server for processing). Using a homebuilt script further allowed us to change parameters like optical sectioning factor, scaling factor and filter frequencies more freely while knowing exactly what impact each had on the final image. The main speed limitation of widefield acquisition on the Mesolens was not the camera exposure but rather handling of the data itself. At full FOV, camera images are approximately 500 Mb large. Each image needed to be transferred from the camera buffer to the PC’s hard drive before a new image could be acquired. As a direct result, reducing acquisition time by a factor of five (1000 ms to 200 ms) only reduced acquisition time for a 25-image z-stack from 32 minutes to 27 minutes (~15% time reduction). The chip shifting mechanism of the VPN-29MC was necessary to obtain Nyquist sampled images but shifting the detector chip through nine positions meant that exposure time for one frame was 1800 ms (9 times 200ms) at minimum. The fastest practical exposure setting was 200 ms, as below this exposure time the readout of the CCD detector array increased frame time to 200ms regardless (according to VPN-29MC user manual). A single detector array of sufficiently small pixel size would potentially increase acquisition time but would not get around the data handling constrains. Furthermore, no such detector with such high pixel number and sensor size compatible with the Mesolens is commercially available at present and custom-built detectors would come at very high cost.

Despite these drawbacks, HiLo mesoscopy is an excellent alternative to CLSM for the Mesolens and offers some advantage over SPIM. Unlike SPIM, HiLo obtains uniform section thickness over the full FOV comparable in sectioning strength to CLSM. The setup for HiLo is considerably simpler with just a diffuser in an otherwise ordinary transmission illumination setup. SPIM would require side illumination, potentially casting lateral shadows across the FOV at optical section thickness upwards of 20 μm for standard Gaussian beams. SPIM also requires specialised sample chambers whereas HiLo can operate with the same sample preparation procedures already in place for the Mesolens’ other imaging modalities.

It has been discussed in detail ^18^ that the strong out-of-focus background in thick samples presents a problem for HiLo imaging by reducing in-focus speckle contrast and hence making it more difficult to distinguish in-focus from out-of-focus regions in the image. To maintain high speckle contrast, a coarser speckle illumination pattern was used here. Rather than decreasing the illumination NA, we chose to implement a variable beam expander in the illumination beam path. By changing the beam diameter illuminating the diffuser, we could adjust the speckle coarseness to suit the sample and maintain sectioning capability, albeit with thicker sections. Although the sections were thicker than the axial resolution, approximately 10 μm for σ=3, sectioning was still superior to what SPIM is capable of on this FOV. The above-mentioned contrast issue affects axial resolution only since lateral resolution is determined by the microscope system.

## Methods

### Theoretical background of Hilo microscopy

HiLo microscopy was described in detail by its developers Lim and Mertz elsewhere ^14,17^ so we repeat only the basic principle of its operation here. HiLo microscopy performs optical sectioning of fluorescent samples by segmenting the image using a weighting function obtained by contrast evaluation of the difference image of a structured illumination image and a uniform illumination image. The uniform image iu is a simple widefield fluorescent image. To obtain the structured illumination image is, the sample is illuminated by a random laser speckle pattern. The in-focus high spatial frequencies of the image are obtained by simply applying a gaussian high-pass filter to a Fourier transformed uniform image such that:

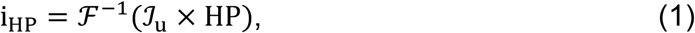

where i_HP_ is the high-pass filtered uniform image, 𝓕 ^−1^ is the inverse Fourier Transform, 𝓘 _u_ is the Fourier Transform of iu and HP is a gaussian high-pass filter with cut-off frequency kc, such that HP(k_c_) = 1/2.

The high spatial frequencies are inherently in focus and thus do not need to be further processed. To obtain the in-focus low spatial frequencies, first the difference image, id, must be calculated

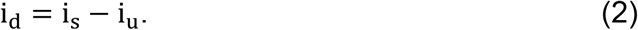

Subtracting i_u_ from i_s_ removes the sample induced bias and allows the evaluation of local speckle contrast to be performed on the variations of the speckle pattern only.

The local contrast of speckle grains tends to zero with defocus and thus allows to distinguish between in-focus and out-of-focus signal. This decay to zero can be accelerated by applying a bandpass filter to id prior to contrast evaluation

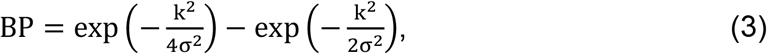

where BP is the bandpass filter, generated by subtracting two Gaussian lowpass filters, k is the spatial frequency and σ is the bandpass filter standard deviation.

Correct evaluation of local speckle contrast is key to separate in-focus from out-of-focus signal. Local contrast evaluation can be performed by calculating the quotient of standard deviation and mean in a local neighbourhood with a sliding window ^18,19^.

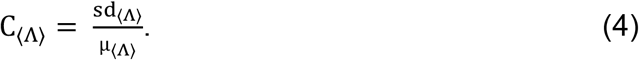

where C_⟨Λ⟩_ is the contrast in the local neighbourhood, evaluated within a sliding window of side length Λ (pixels). Sd_⟨Λ⟩_ and μ_⟨Λ⟩_ are the standard deviation and mean intensity in the local neighbourhood respectively.

The side length Λ of the sliding window is determined depending on the cut-off frequency kc as described in reference ^18^.

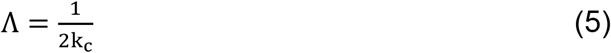

Applying the local contrast as a weighting function to i_u_ results in a coarse image of in-focus low spatial frequencies.

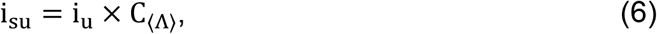

where i_su_ is the weighted uniform illumination image. By applying a gaussian low-pass filter LP complementary to HP, i.e. LP + HP = 1, the in-focus low spatial frequencies are obtained

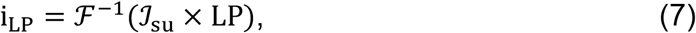

where i_LP_ is the in-focus low spatial frequency image, 𝓘 _su_ is the Fourier Transform of i_su_ and LP is the complementary low-pass filter. To ensure a smooth transition between i_LP_ and i_HP_, a scaling factor is calculated

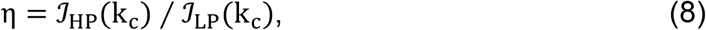

where η is the scaling factor, 𝓘 _HP_(k_c_) and 𝓘 _LP_(k_c_) are the Fourier Transforms of i_HP_ and i_LP_ respectively evaluated at the cut-off frequency k_c_.

The final optically sectioned HiLo image is obtained by adding the in-focus high and low spatial frequency images together.

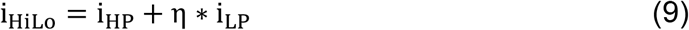

Where i_HiLo_ is the final optically sectioned image. By setting k_c_=0.18σ ^17,18^, the optical sectioning strength can be controlled by changing only the σ parameter.

A schematic of the setup is shown in **Figure 3**. A Coherent Sapphire 488-10 CDRH laser was used as a light source. The beam was guided through a beam expander (Thorlabs BE02-05, 2x-5x variable zoom Galilean beam expander), increasing the beam diameter from 1 mm to 2-5 mm. Subsequently the beam illuminated a 1500 grit ground glass diffuser (DG20-1500, Thorlabs). The diffuser was glued to a DC motor controlled via an Arduino Uno board connected to a PC via USB. It was imaged onto the back aperture of the 0.6 NA Mesolens condenser (Mesolens Ltd.) using an aspheric lens with 0.6 NA (ACL5040U-A, Thorlabs). With the diffuser stationary, a speckle pattern was generated in the sample. Rotating the diffuser via the DC motor (6/9 V, 12000±15% rpm) resulted in uniform illumination, thus allowing acquisition of both uniform and speckle illumination images in quick succession about, 1 minute raw acquisition time per image pair on the full 4.4 mm FOV of the camera. Images were acquired with a thermoelectric Peltier cooled camera (VNP-29MC, Vieworks) with a chip-shifting mechanism. The chip-shifting mechanism was essential to benefit from the large FOV and high resolution (700 nm lateral, 7 μm axial ^1^) provided by the Mesolens. The camera could be operated without chip-shift at a resolution of 6576 × 4384 pixels (28.8 Megapixel), with 4x chip-shift at 13152 × 8768 pixels (115.3 Megapixel) and with 9x chip-shift at 19728 × 13152 pixels (259.5 Megapixel). For HiLo imaging with the Mesolens, the chosen mode was always 9x chip-shift. In this mode, the sampling rate was 4.456 px/μm, corresponding to a 224 nm pixel size, satisfying Nyquist sampling. The minimum frame time of the Stemmer was 200 ms resulting in acquisition time for one full FOV image with 9x pixel shift of 1800 ms excluding time to transfer the image data from the camera to the PC which usually took on the order of 10 seconds. In practice, acquisition of one image took ~12-15 seconds including transfer of data and beginning of new image capture.

**Figure 3:**
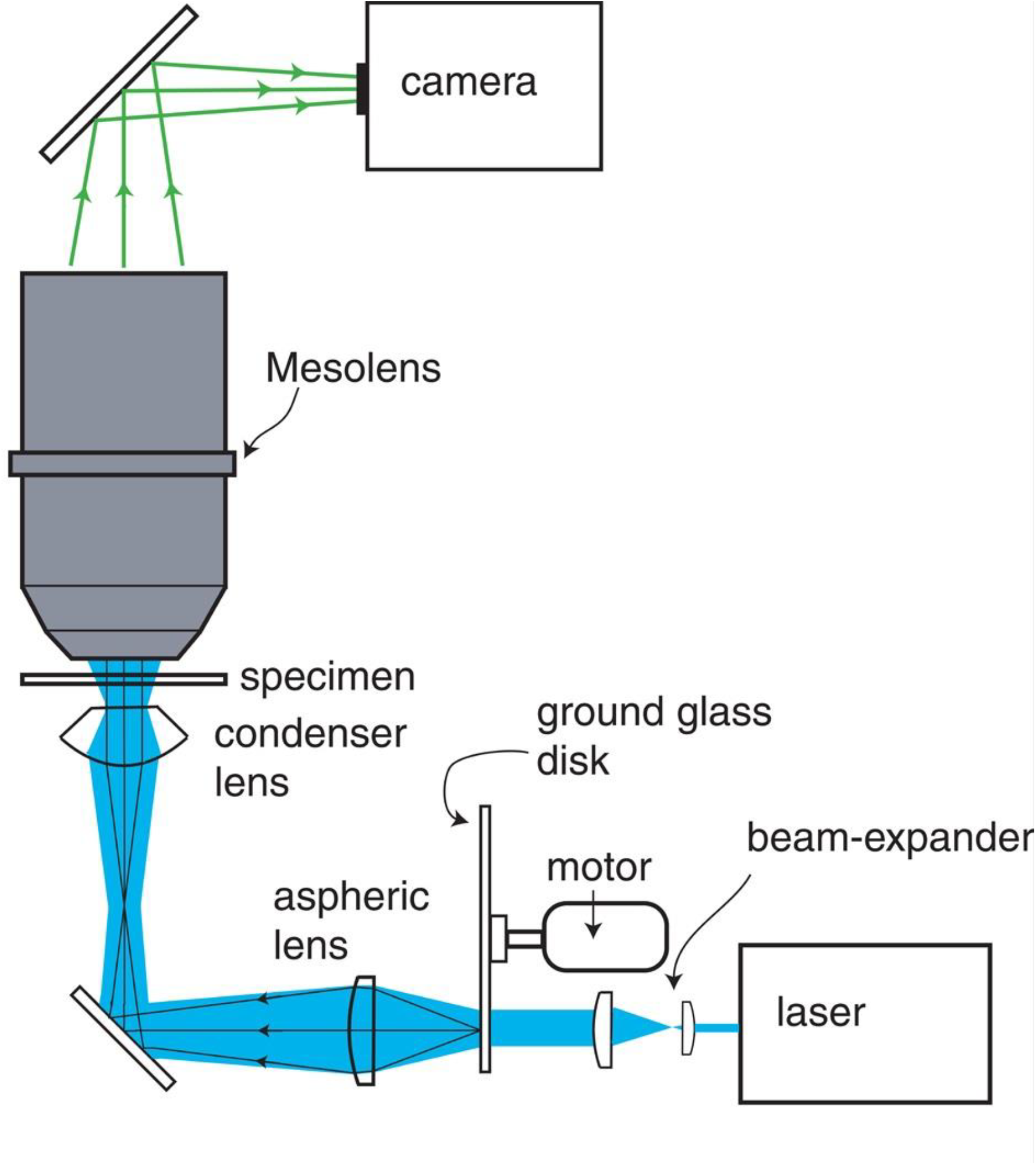
Experimental setup to generate speckle and uniform illumination. The laser emitted light at 488 nm. The beam was expanded to a maximum diameter of approximately 5 mm and illuminated the 1500 grit ground glass diffuser. The diffuser was imaged by the 0.6 NA aspheric lens onto the back aperture of the Mesolens condenser. A stationary diffuser resulted in speckle illumination of the sample while rotating diffuser illuminated the sample uniformly. The sample was then imaged by the Mesolens onto a camera detector. Not shown in this diagram are two mirrors that are placed before and after the beam expander to guide the laser beam.

### Data processing

To process the speckle and uniform images a MATLAB (R2016b version 9.1.0.441655, 64bit, MathWorks, Inc.) script was written that performed HiLo imaging in the same manner as described in the previous section. This allowed more control over individual parameters (optical sectioning factor, low frequency scaling and cut-off frequency) and opened the possibility to use the parallel processing toolbox of MATLAB to use a graphics processing unit (GPU) for processing (the script will be published on the Github repository). Because of the file size of Mesolens images, it was necessary to process z-stacks of samples on a server as commercially available desktop PCs do not have sufficient memory to open or process such large files. The server was a Dell PowerEdge R740 with 1TB RAM and an NVIDIA Quadro P4000 GPU with 8GB video memory.

### Measuring the optical sectioning capability of the HiLo mesoscope

To determine the optical sectioning strength of the HiLo method, a thin fluorescent layer was set at a tilt and then imaged such that the resulting image would show the fluorescent layer coming into focus in the centre of the field of view and go out of focus towards the left and right. This method was adapted after ^20^. To prepare the thin fluorescent layer, first a 170 μm thick microscope cover slip (22 mm × 22 mm, #1.5, Thermo Fisher Scientific) was rinsed in dry acetone (Acetone 20066.330, VWR Chemicals). It was then submerged in an APTMS-acetone (3- Aminopropyldrimethoxysaline, 281778-100ML, Sigma Aldrich) solution for six hours (0.2 mL APTMS, 9.8 mL of dry acetone). After this period, the cover slip was rinsed three times in dry acetone and blow-dried with compressed air. The cover slip was put in a 10 μM solution of fluorescein salt (Fluorescein sodium salt, 46960-25G-F, Sigma Aldrich) in distilled water. Care was taken to only let one side of the cover slip get in contact with the fluorescein solution to avoid having two thin fluorescent layers (one on either side). The bath was carefully wrapped in aluminium foil and left overnight in a dark place. The next day, the cover slip was rinsed with distilled water twice and again blow-dried with compressed air. Finally, the cover slip was mounted on a microscope slide with the dye-coated surface in contact with the slide and was sealed with nail varnish. Imaging was done with glycerol immersion.

### Mouse hippocampal neuron sample preparation

The mouse hippocampal neuron sample was prepared from C57BL/6J pups (1-2 days old) as described previously 21,22 and fluorescently stained 23,24. All experimental procedures were performed in accordance with UK legislation including the Animals (Scientific Procedures) Act 1986 and with approval of the University of Strathclyde Animal Welfare and Ethical Review Body (AWERB). In short, neurons were fixed in ice-cold 4% paraformaldehyde (PFA). The sample was then incubated with a primary anti-mouse antibody (anti-βIII-tubulin (1:500), Sigma-Aldrich) and fluorescently labelled using a secondary antibody (anti-rabbit AlexaFluor 488 (1:200), Thermo Fisher Scientific). The fixed and stained sample was mounted onto a glass microscope slide (VWR, UK) using Vectashield mounting medium (H-1200, Vector Laboratories) and imaging was performed with glycerol immersion on the Mesolens.

### 5-day-old zebrafish larva sample preparation

The zebrafish were fixed in ethanol: glacial acetic acid at a 3:1 ratio at 4°C for several days, rehydrated through an ethanol/water series and stained with 0.001% Acridine Orange (A1301, Thermo Fisher Scientific) in phosphate-buffered saline, washed in PBS, dehydrated in an ethanol series and cleared in xylene before mounting in Fluoromount ^25^ (Fluoromount is no longer available commercially: we would advise Histomount (Thermo Fisher Scientific) as similar substitute). Imaging was performed with glycerol immersion on the Mesolens.

## Acknowledgement

We thank Dr David Li and Ross Scrimgeour for their help with digital image processing.

This work was jointly funded by the University of Strathclyde, Glasgow and the Hong Kong University of Science and Technology (HKUST), Hong Kong.

This work was supported by the Medical Research Council [MR/K015583/1].

## Author contribution statement

G.M. conceived the experiments, J.S. performed the experimental work and data analysis, A.F. and T.B. prepared the mouse hippocampal neuron sample, W.B.A. prepared the zebrafish specimen, J.D. wrote the image acquisition software based on WinFluor. All authors reviewed the manuscript.

## Additional information

The authors declare no competing financial and/or non-financial interest.

## Supplementary Data

**Supplementary Video 1: Acridine Orange stained zebrafish.** The first part of the movie shows a cropped FOV (smaller than the Mesolens full FOV but large enough to see the whole zebrafish) 61- image z-series (at 3 μm z-steps) of a zebrafish. The second part of the movie shows a software zoom to the region where the eye is located and the third part is, again, a z-series of the ROI. In the third part, individual cells are clearly visible and the depth structure of the sample becomes obvious. The sample was exited at a wavelength of 488 nm and fluorescence detected at 540nm as before. The optical sectioning paramter σ was set to 3 for these data with coarser speckle illumination pattern to avoid speckle structure artefacts in the final image.

**Supplementary Video 2: HiLo imaging of fixed and fluorescently stained mouse hippocampal neurons**. The first part of the video shows HiLo processed 25-image z-series (at 3 μm z-steps) of fixed and stained hippocampal mouse neurons obtained with the full FOV of the Mesolens. Raw data has been contrast adjusted for presentation and the Fire LUT has been applied to increase contrast further. The second part of the movie is a zoom into a small region of interest (ROI) with only a few neurons in the FOV. The third part of the movie is a z-series of that ROI. The detail shown in the third part of the video is representative of the detail of the entire FOV without the need to re-acquire data. The zoom is purely digital. The specimen was excited at a wavelength of 488 nm and fluorescence was detected by the camera at an emission wavelength peak of 540 nm with 20 nm full-width-half-maximum spectral bandwidth. The sectioning parameter σ was set to 1 for these data. The apparent sweep of brightness across the specimen is a result of it not being flat relative to the direction of illumination.

## References

1. McConnell, G. et al. A novel optical microscope for imaging large embryos and tissue volumes with sub-cellular resolution throughout. Elife 5, 1–15 (2016).

2. McConnell, G. & Amos, W. B. Application of the Mesolens for subcellular resolution imaging of intact larval and whole adult Drosophila. Preprint at https://www.biorxiv.org/content/early/2018/02/19/267823 (2018).

3. Hwang, J. Y. et al. Multimodal wide-field two-photon excitation imaging: characterization of the technique for in vivo applications. Biomed. Opt. Express 2, 356 (2011).

4. Amor, R. et al. Widefield Two-Photon Excitation without Scanning: Live Cell Microscopy with High Time Resolution and Low Photo-Bleaching. PLoS One 11, e0147115 (2016).

5. Yew, E. Y. S., Sheppard, C. J. R. & So, P. T. C. Temporally focused wide-field two-photon microscopy: Paraxial to vectorial. Opt. Express 21, 12951 (2013).

6. Oron, D., Tal, E. & Silberberg, Y. Scanningless depth-resolved microscopy. Opt. Express 13, 1468 (2005).

7. Siedentopf, H. & Zsigmondy, R. Über Sichtbarmachung und Größenbestimmung ultramikoskopischer Teilchen, mit besonderer Anwendung auf Goldrubingläser. Ann. Phys. 315, 1–39 (1902).

8. Voie, A. H., Burns, D. H. & Spelman, F. A. Orthogonal-plane fluorescence optical sectioning: Three-dimensional imaging of macroscopic biological specimens. J. Microsc. 170, 229–236 (1993).

9. Huisken, J. & Stainier, D. Y. R. Even fluorescence excitation by multidirectional selective plane illumination microscopy (mSPIM). Opt. Lett. 32, 2608 (2007).

10. Yang, Z. et al. A compact Airy beam light sheet microscope with a tilted cylindrical lens. Biomed. Opt. Express 5, 3434–42 (2014).

11. Vettenburg, T. et al. Light-sheet microscopy using an Airy beam. Nat. Methods 11, 541–544 (2014).

12. Gustafsson, M. G. L. Surpassing the lateral resolution limit by a factor of two using structured illumination microscopy. J. Microsc. 198, 82–87 (2000).

13. Gustafsson, M. G. L., Agard, D. A. & Sedat, J. W. Method and apparatus for three-dimensional microscopy with enhanced depth resolution. 23–25 (1997).

14. Lim, D., Chu, K. K. & Mertz, J. Wide-field fluorescence sectioning with hybrid speckle and uniform-illumination microscopy. Opt. Lett. 33, 1819–1821 (2008).

15. Ströhl, F. & Kaminski, C. F. Speed limits of structured illumination microscopy. Opt. Lett. 42, 2511 (2017).

16. Orieux, F., Sepulveda, E., Loriette, V., Dubertret, B. & Olivo-Marin, J. C. Bayesian estimation for optimized structured illumination microscopy. IEEE Trans. Image Process. 21, 601–614 (2012).

17. Lim, D., Ford, T. N., Chu, K. K. & Mertz, J. Optically sectioned in vivo imaging with speckle illumination HiLo microscopy. J. Biomed. Opt. 16, 016014 (2011).

18. Mazzaferri, J. et al. Analyzing speckle contrast for HiLo microscopy optimization. Opt. Express 19, 14508 (2011).

19. Duncan, D. D., Kirkpatrick, S. J. & Wang, R. K. Statistics of local speckle contrast. J. Opt. Soc. Am. A 25, 9 (2008).

20. B. Amos, G. McConnell, T. W. Confocal Microscopy. in Comprehensive Biophysics (ed. Egelman, E. H.) 3–23 (Elsevier B.V., 2011).

21. McNair, K., Davies, C. H. & Cobb, S. R. Plasticity-related regulation of the hippocampal proteome. Eur. J. Neurosci. 23, 575–580 (2006).

22. Gan, J., Greenwood, S. M., Cobb, S. R. & Bushell, T. J. Indirect modulation of neuronal excitability and synaptic transmission in the hippocampus by activation of proteinase-activated receptor-2. Br. J. Pharmacol. 163, 984–994 (2011).

23. Ritchie, L. et al. Toll-like receptor 3 activation impairs excitability and synaptic activity via TRIF signalling in immature rat and human neurons. Neuropharmacology 135, 1–10 (2018).

24. Abdul Rahman, N. Z. et al. Mitogen-Activated Protein Kinase Phosphatase-2 Deletion Impairs Synaptic Plasticity and Hippocampal-Dependent Memory. J. Neurosci. 36, 2348–2354 (2016).

25. Gurr, E. Fluorescence microscopy. J. R. Nav. Med. Serv. 37, 133–40 (1951).

